# *Trichodesmium Erythraeum* produces a higher photocurrent than other cyanobacterial species in bio-photo electrochemical cells

**DOI:** 10.1101/2022.01.23.477402

**Authors:** Yaniv Shlosberg, Dina Spungin, Neta Holland, Gadi Schuster, Ilana-Berman Frank, Noam Adir

## Abstract

In recent years, the increase in world energy consumption, and the worries from potential future disasters that may derive from climate change have inspired the motivation to develop renewable energy technologies. One of the promising methods is the utilization of whole bacterial cells to produce photocurrent in a bio-photo electrochemical cell (BPEC). The photocurrent derives from the photosynthesis pathway, while the redox couple NADP^+^/NADPH perform cyclic electron mediation between photosystem I inside the cells, and the anode. Over the years, various cyanobacterial species were utilized in diverse BPECs setups, while the photocurrent was enhanced by the addition of natural electron mediators such as NAD^+^, NADP^+^, Cytochrome C, Vitamin B1, and the artificial mediator potassium ferricyanide. The cyanobacterium *Trichodesmium Erythraeum* (*Te*) is a marine species that consist of high content of Phycocyanin and Phycoerythrin pigments that play a major role in photosynthesis enhancement. In this work, we produce for the first-time photocurrent from *Te*. We apply 2D-fluorescence measurements to detect its NADPH secretion and show that its photocurrent production is enhanced as a function of increasing electrolyte salinity. Finally, we produce photocurrent from additional cyanobacterial species: *Synechocystis* sp. PCC6803, *Synechococcus elongatus PCC 7942, Acaryochloris marina* MBIC 11017, and *Spirulina*, using their cultivation medium as electrolytes in the BPEC. We show that TE produces a photocurrent intensity that is significantly greater than all other species with and without the addition of exogenous electron mediators. The utilization of TE may pave the way toward the establishment of marine clean energy technologies.

## Introduction

In recent years, the predictions for disasters that may occur in the future because of climate change have motivated scientists around the world to develop new clean energy technologies. One of the promising approaches that were invented a few decades ago, was the invention of microbial fuel cells (MFCs). ^1–9^ This method is based on the ability of bacterial species to conduct exo-electrogenic activity and reduce the anode in an electrochemical cell to produce electrical current. The current production can be done by 2 different mechanisms: direct electron transport (DET), and mediated electron transport (MET). The DET mechanism is performed by proteins complexes such as *pili* and MTR that are capable of charge transfer. ^10–15^ Among the most efficient species that can conduct DET are the bacteria *Shewanella oneidensis* and *Geobacter sulfurreducens* that are rich in these kinds of complexes. ^10–15^ MET can be done by the addition of an exogenous electron mediator that can transfer electrons between the cells and the anode. ^6,7,16–19^ Over the years, various electron mediators were reported including potassium ferricyanide (Fe(CN)_6_), quinones, and flavines and phenazines derivatives. ^4,20–24^

To maintain the viability of bacteria in MFCs over time, sugar sources must be supplied to the cells. One of the solutions for this is the integration of MFCs systems in the soil together with plants’ roots. ^25–27^ The plants conduct photosynthesis and secrete some of the formed sugar through their roots, while these sugar molecules can feed the bacteria. The utilization of plants as a direct sugar source is low cost and considered to be eco-friendly because they absorb CO_2_ from the atmosphere.

Another eco-friendly energy solution that was evolved from MFCs is the bio-photo electrochemical cell. This approach integrates whole photosynthetic organisms such as plants’ green tissues, ^28^ seaweeds^29^, microalgae^30^, and cyanobacteria^31–38^ with bio-electrochemical cells to produce photocurrent. The electron source in BPECs originates from the respiration and photosynthetic pathway.^33,34^ Some of the NADPH molecules that are formed in photosynthesis can exit the organisms and donate electrons to the anode of the BPEC to produce electrical current. ^33^ The cultivation of photosynthetic organisms is simpler and chipper than non-photosynthetic bacteria since they can synthesize their carbon source by doing photosynthesis. ^39^ Enhancement of the photocurrent can be achieved by exogenous addition of the native electron mediator NADP^+^, and other bio-compatible electron mediators such the metabolites NAD^+^ and vitamin B1, or the chemical Fe(CN)_6_. ^29,30,33,34^ In microorganismal-based BPECs, the addition of the redox-active metabolites enhance the photocurrent by ∼ 6 times, ^33^ while Fe(CN)_6_ is more efficient enhancing it by ∼ 20 times. ^30^

A major important factor that significantly enhances the current production in BPEC is the salinity of the electrolyte that induces its conductivity. ^29^ However, unlike non-biological electrochemical cells that are mostly not sensitive to high salt concentration, the environment of BPECs must be compatible to enable the survival of the used organisms. The utilization of marine organisms is therefore a big advantage as they can tolerate high electrolyte salinity. ^29,30^

One of the most abundant marine cyanobacterium is *Trichodesmium Erythraeum* (*Te)*.

In this work, we use for the first time *Te* to produce photocurrent in a BPEC. We apply 2D-fluorescence measurements to show that the secretion of NAD(P)H by *Te* is enhanced upon solar illumination. We contrast the ability of *Te* vs. 4 other cyanobacterial species: *Acaryochloris marina* MBIC 11017 (*Am*), *Synechocystis* sp. PCC 6803 (Syn), and *Synechococcus* elongatus PCC 7942 (Se), and *Spirulina* (Spi) to produce photocurrent in a BPEC using their cultivation media as electrolytes. We show that *Te* significantly produces more photocurrent than the other species without and with the addition of the exogenous electron mediators: NADP^+^, vitamin B1, Cytochrome C, and Fe(CN)_6_.

## Results

### Photocurrent production of TE at increasing salinity

In our previous works, we showed that the salinity of the electrolyte plays a key role in the enhancement of the current production in micro, and macroalgal-based BPECs. For this reason, marine photosynthetic organisms that habitat may be good candidates to be utilized in BPECs, since they can tolerate high salinity. The measurements setup for photocurrent production is the same setup that was described in our previous work ^30,33^. The setup consisted of screen-printed electrodes (SPEs) with a graphite anode, a platinum cathode, and silver coated with a silver chloride reference electrode. Unlike most cyanobacterial solutions that form a homogenous rigid suspension, TE filaments tend to float and mostly form a heterogenous solution in which they do not stick to each other. Also, they are sensitive to centrifugation that might damage their viability. To cover the entire surface area of the anode, a concentrated solution of TE was added into a small plastic cylinder with a diameter of ∼ 0.4 cm^2^ (the diameter of the anode) that was placed on top of the anode (fig.1a). After ∼10 min the cells sunk and covered the anode. Chronoamperometry (CA) of TE cells was measured (Fig.1) with an applied bias of 0.5 V on the anode. Solar illumination 1.5 SUN (1500W / m^2^) was performed applied from above the cell by a Solar simulator.

**Fig.1.**
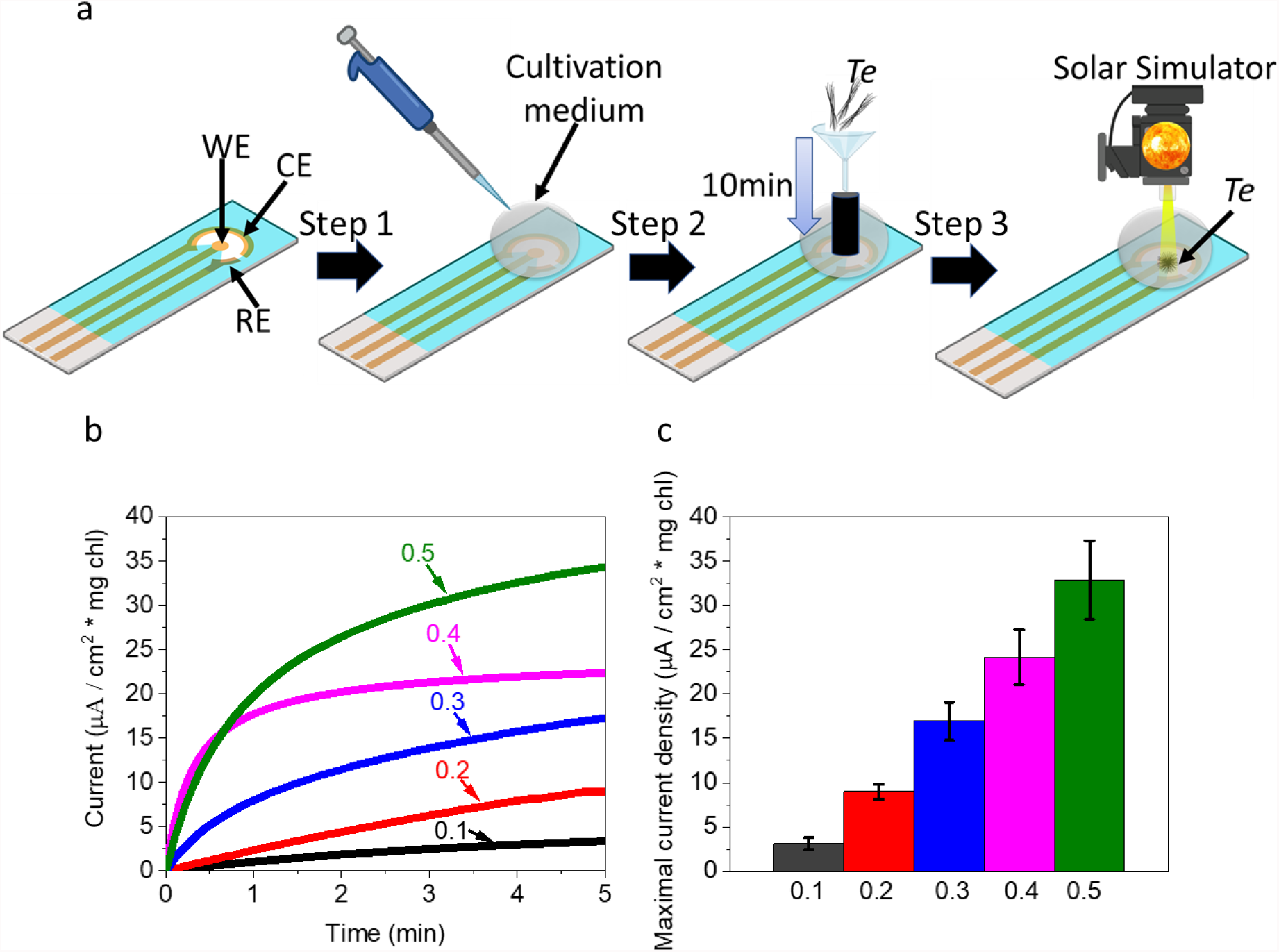
Photocurrent production of TE at increasing salinity. CA of *Te* cells was measured in an electrolyte solution with increasing NaCl concentrations (0.1 – 0.5 M). **a** A description of the procedure for measuring the photocurrent production of *Te* using screen-printed electrodes. Step 1 – A drop of 100 µL electrolyte is added to cover the entire area of the electrodes. Step 2 – A small plastic cylinder with the diameter of the working electrode (WE) is placed on it inside the drop. concentrated *Te* cells are added into the cylinder to let them precipitate for ∼ 10 min. Step 3 – the cylinder is gently removed letting the *Te* cells cover the entire surface of the WE. Illumination of the *Te* is conducted by a solar simulator that is placed vertically to the WE. The solar light intensity was 1000 W / m^2^. **b** Representative CA measurements of *Te* in light in electrolytes with increasing NaCl solutions: 0.1, 0.2, 0.3, 0.4, and 0.5 M. **c** Maximal photocurrent densities of *Te* in the electrolytes with increasing NaCl solutions. The error bars represent 3 the standard deviation over 3 independent measurements.

### NADPH is the major electron mediator in TE-based BPECs

In our previous work, ^33^ we showed that upon association of various cyanobacterial species with the anode of a BPEC, they secrete NADH and NADPH into the external cellular solution (ECS), while NADPH is the major electron mediator that is responsible for the photocurrent production. We wished to assess whether the same electron transport mechanism occurs in TE-based BPECs. To explore this, CA of TE cells was measured in dark and light for 10 min, right after the measurement, the cells were filtrated through a 0.22µM PVDF and 2D-fluorescence maps of their ECS was measured. As previously reported for other cyanobacterial species^33^, the fluorescent spectral fingerprint of NAD(P)H was obtained with a maximal intensity around (λ_Ex_= 350 nm, λ_Ex_= 450 nm), while its intensity was ∼ 10 times higher in light than in dark. (Fig. 2a,b). The intensity difference is ∼ 2 times bigger than previously obtained for other cyanobacterial species^33^. Quantification of the NAD(P)H was performed based on a calibration curve of increasing concentrations of NADH vs. their fluorescence intensity at (λ_Ex_= 350 nm, λ_Ex_= 450 nm). (Fig. 2c)

**Fig. 2.**
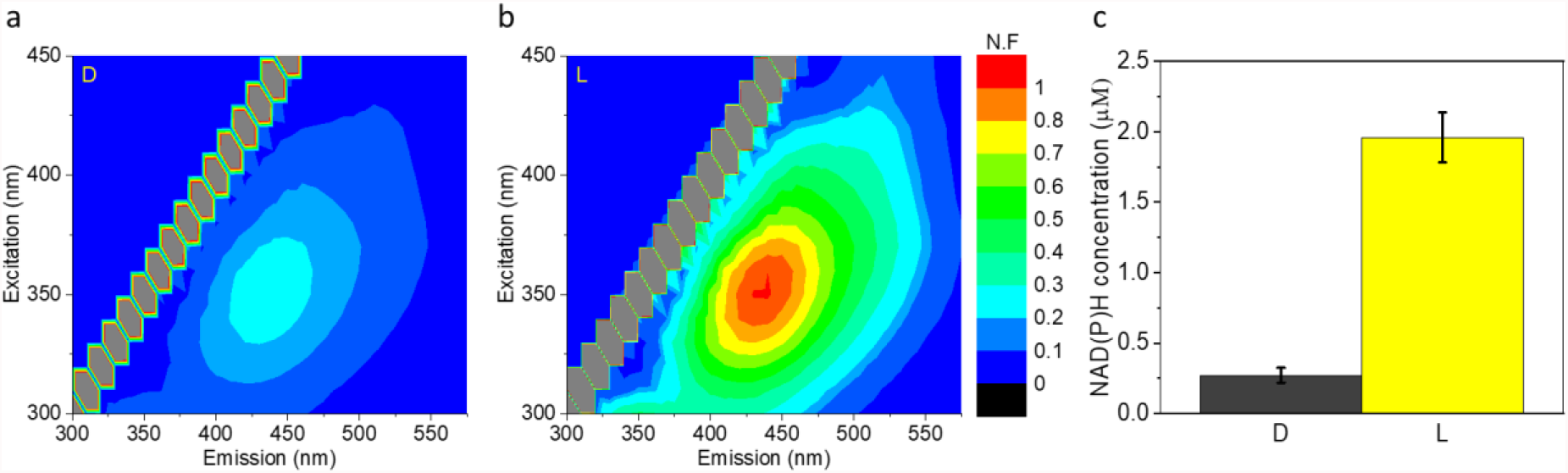
NADPH is the major electron mediator in TE-based BPECs. The NAD(P)H concentrations in the ECS of *Te* after CA measurements in dark and light were estimated based on 2D-FM measurements. **a** 2D-FM of *Te* ECS after CA in dark. **b** 2D-FM of *Te* ECS after CA in light. **c** The estimated concentration of NAD(P)H in the ECS of *Te* after CA in dark and light. The error bars represent the standard deviation over 3 independent repetitions.

### External electron transport of cyanobacteria in dark

Based on the observation that *Te* secretes NAD(P)H molecules also in dark, we wished to assess whether intact *Te* and the cyanobacterial species *Trychodesmium Erythraeum* (*Te*), *Acaryochloris marina MBIC 11017* (*Am*), the freshwater species *Synechocystis sp. PCC 6803* (*Syn*), and *Synechococcus elongatus* PCC 7942 (*Se)*, and *Spirulina* (*Spi*) produce an electrical current in a BPEC in the absence of light. CA of the 5 species was measured using their own cultivation media as electrolytes. (Fig. 3). No significant current was obtained for any of the species except for *Spi*, which produced a maximal current density of ∼ 8 µA / cm^2^ * mg Chl. (Fig. 3) We postulated that this current may derive from the hydroxide ions in the cultivation media of *Spi* that may reduce the anode, as previously reported for alkaline-based electrochemical cells. ^40^ To elucidate this, CA in dark was measured for the cultivation media of *Spi* (without the cells) and a basic water solution (the pH value of the 2 solutions was 9). In both measurements, the maximal obtained current production was the same as obtained in the measurements with the *Spi* cells (Fig. 3), indicating that the cells do not produce any significant current in dark.

**Fig. 3.**
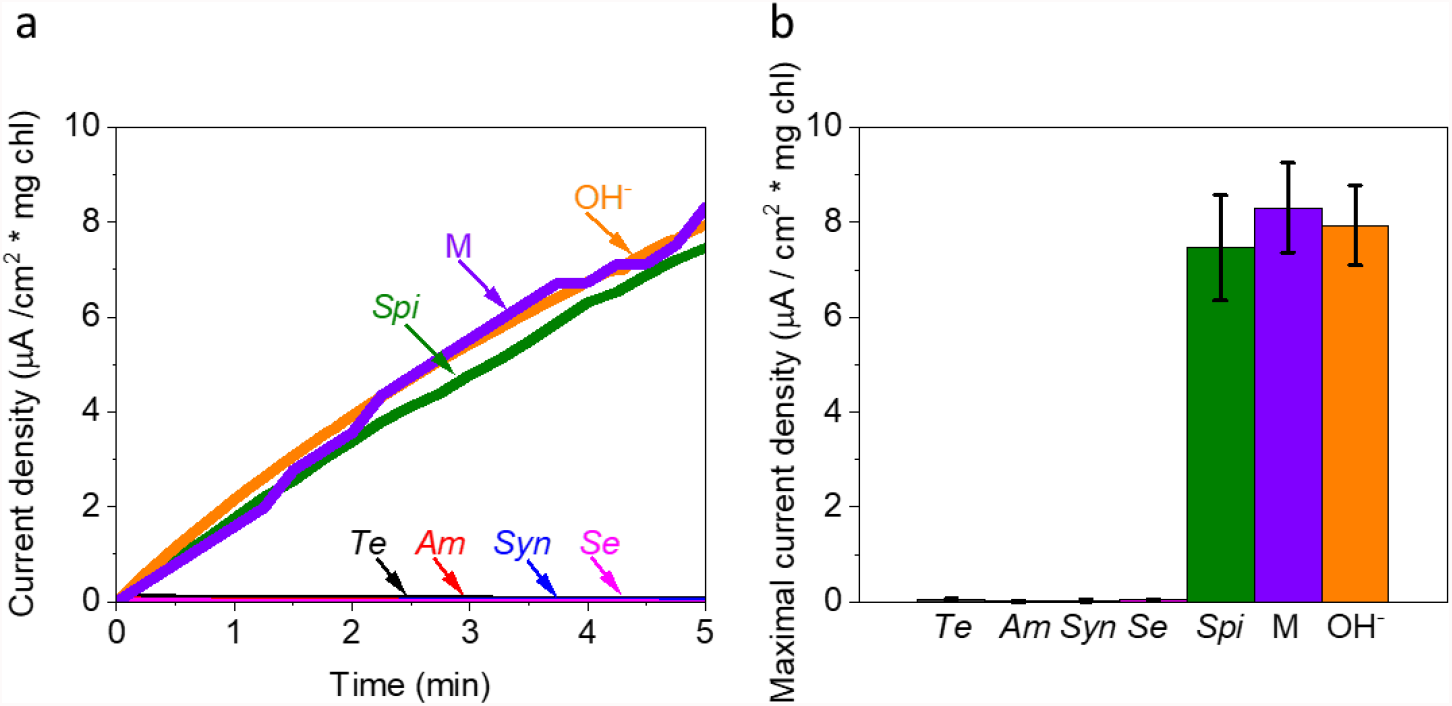
External electron transport of cyanobacteria in dark. CA of the 5 cyanobacterial species: *Te, Am, Syn, Se, Spi*, the cultivation media of *Spi* (pH = 9), and a basic water solution (pH = 9), were measured in dark. **a** Representative CA measurements of *Te* (black), *Am* (red), *Syn* (blue), *Se* (magenta), *Spi* (green), the cultivation media of *Spi* (pH = 9) (M, purple), and a basic water solution (pH = 9) (OH^-^, orange). **b** Maximal current densities. The error bars represent the standard deviation over 3 independent repetitions. The current density in the measurements of the *Spi* cultivation media and the basic solutions were normalized to the same Chl concentrations that were obtained for the *Spi* cells (∼ 0.1 mg).

### The photocurrent in cyanobacterial BPECs derives from the respiration pathway and PSI

In our previous works^33,34^, we showed that the addition of the PSII inhibitor the herbicide 3-(3,4-dichlorophenyl)-1,1-dimethylurea (DCMU), or Glucose enhance the photocurrent production in BPECs that are based on *Syn*. We wished to explore if this photocurrent enhancement also occurs for other cyanobacterial species. To do that, CA of *Te, Am, Syn, Se*, and *Spi* was measured in light with addition of 100 µM DCMU right before the measurement, or after 1 h incubation with 5mM Glucose. The addition of DCMU, and Glucose has enhanced the photocurrent of all 5 species by ∼ 2 and 3 times respectively. (Fig. S1)

### Intact *TE* produces more current than other species

Optimization of a BPEC performance typically involves 2 main systems: the electrochemical setup and the biological component. Optimization of the electrochemical mostly includes the composition of the electrodes, the illumination intensity, the electrolyte, and the applied bias on the working electrode, and the addition of exogenous electron mediators. These are limited by the tolerance of the organisms to the electrochemical environment. A big advantage for using marine organisms derives from their ability to tolerate high salinity, which can significantly enhance the current production when used as the electrolyte of the BPEC^28–30^. We wished to study the ability of various intact cyanobacterial cells to produce photocurrent in their cultivation media (that represents their natural habitat environment). CA of the marine species *Te, Am, Syn, Se*, and *Spi* was performed in light using the cultivation media of the cells as an electrolyte. (Fig. 4) Maximal photocurrents densities of ∼35, 10, 5, 5, and 20 µA / cm^2^ mg Chl were obtained for *Te, Am, Syn, Se*, and *Spi* respectively. The obtained results show that intact cells of *Te* produce a current that is significantly higher than the other species. One of the causes for this may be the high salinity of the cultivation medium of *Te*. Interestingly, the photocurrent that was produced from the marine species *Am* was not significantly greater than obtained for the freshwater cyanobacterial species.

**Fig. 4.**
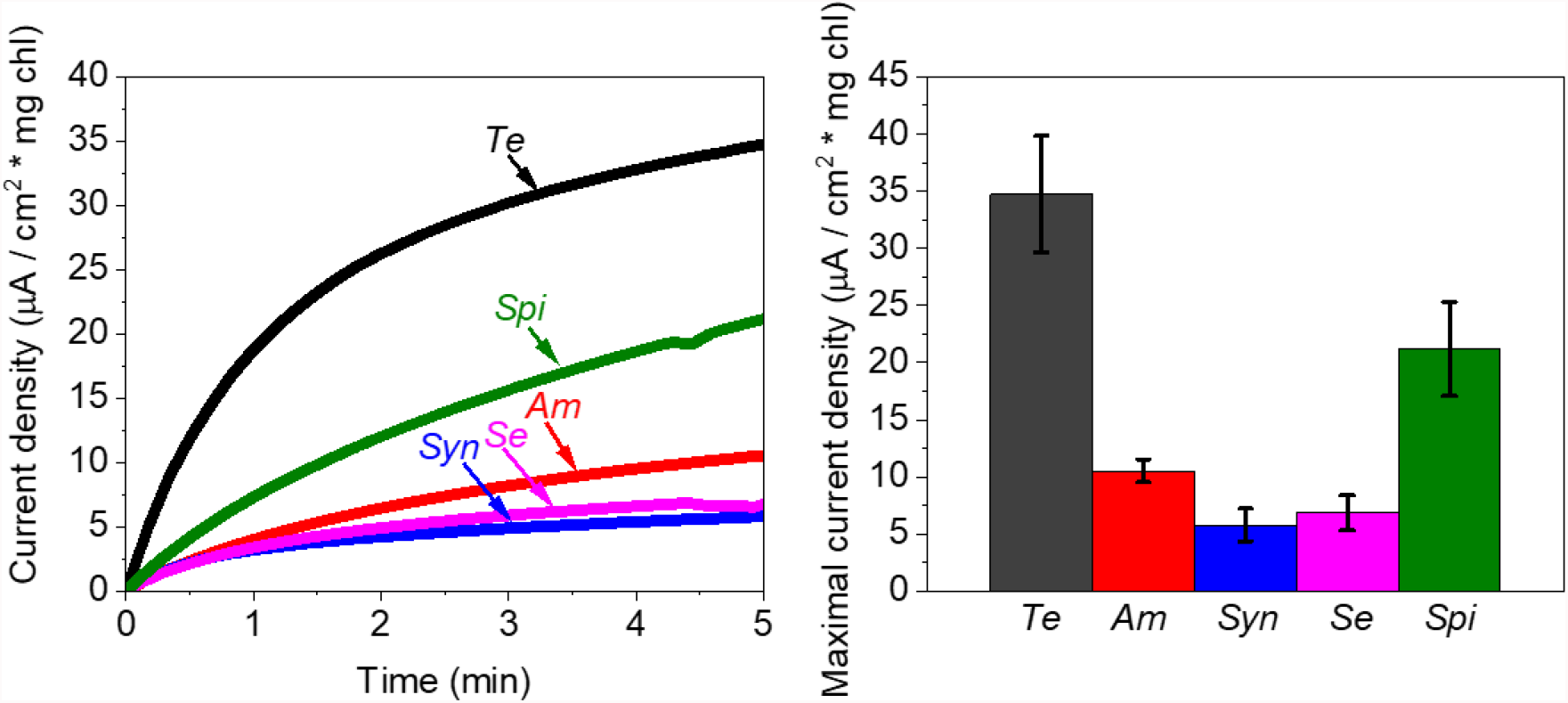
Intact TE produces more current than other species. CA of 5 intact cyanobacterial species in light. **a** Representative CA of *Te* (black), *Am* (red), *Syn* (blue), *Se* (Magenta), and *Spi* (green). **b** Maximal photocurrent densities of *Te* (black), *Am* (red), *Syn* (blue), *Se* (Magenta), and *Spi* (green). The error bars represent the standard deviation over 3 independent measurements.

In nature, the filamentous structure of *Te* that has a low density enables the floatation of the cells to the water surface where the illumination of the sunlight is stronger. It also consists of a relatively high amount of pigment such as phycoerythrin, and phycocyanin that enhances its photosynthesis. In contrast with *Te, Am* habitats in deep water where the sunlight intensity is low. We suggest that illumination at high light that is unnatural for *Am* may initiate photoinhibition mechanisms such as the disconnection of the phycobilisome, which lowers the photosynthesis. This reduction in photosynthesis decreases the formation of photocurrent production. However, the photocurrent is still greater than for the freshwater species, probably because of the high electrolyte salinity of the *Am* cultivation medium.

### TE produces more current than other species in the presence of exogenous electron mediators

In our previous works, we showed the current production of various photosynthetic organisms can be induced by the addition of the exogenous electron mediators NAD^+^, NADP^+^, and potassium ferricyanide. ^29,30,33^ We also showed that photocurrent enhancement can be done in microalgae by the addition of Thiamine (vitamin B1). ^30^ We next wished to explore whether vitamin b1 can also enhance the photocurrent in cyanobacteria and if *TE* produces more current than other cyanobacterial species when exogenous electron mediators are added. To explore this, CA measurements of *Te, Am, Syn, Se*, and *Spi* were measured after 1 h incubation in 5 mM solutions of NADP^+^ and vitamin b1, and without a prior incubation in a 5 mM solution of ferricyanide and Cyt. (Fig. 4) In all measurements, *Te* produced the highest photocurrent Which is about 3 – 6 times greater than the other species.

### A schematic description of the factors that enhance the photocurrent in TE and other cyanobacterial-based BPECs

Based on the results, we suggest a model that describes the electron transfer mechanisms and the unique properties of the different cyanobacterial species that contribute to the current enhancement in the BPEC)Fig. 6). The model shows that *Te* produces a photocurrent that is greater than the other 4 species. In dark, none of the species can produce a current that originates from the cells. However, the media of *Spi* consist of OH^-^ that can reduce the anode to produce current. *Am*-based BPECs can work at high salinity and exploit the benefit of Chl d that broadens the light absorption and enhance the photosynthesis of *Am*. Despite these 2 benefits, *Am* does not produce a greater photocurrent than the freshwater cyanobacterial species *Syn*, and *Se*, probably because of its inability to tolerate high light. For all cyanobacterial species, the photocurrent is enhanced upon utilization of exogenous electron mediators, while their contribution to the photocurrent enhancement of Cyt, B1, NADP^+^, and FeCN_(6)_ is increasing respectively.

**Fig. 5.**
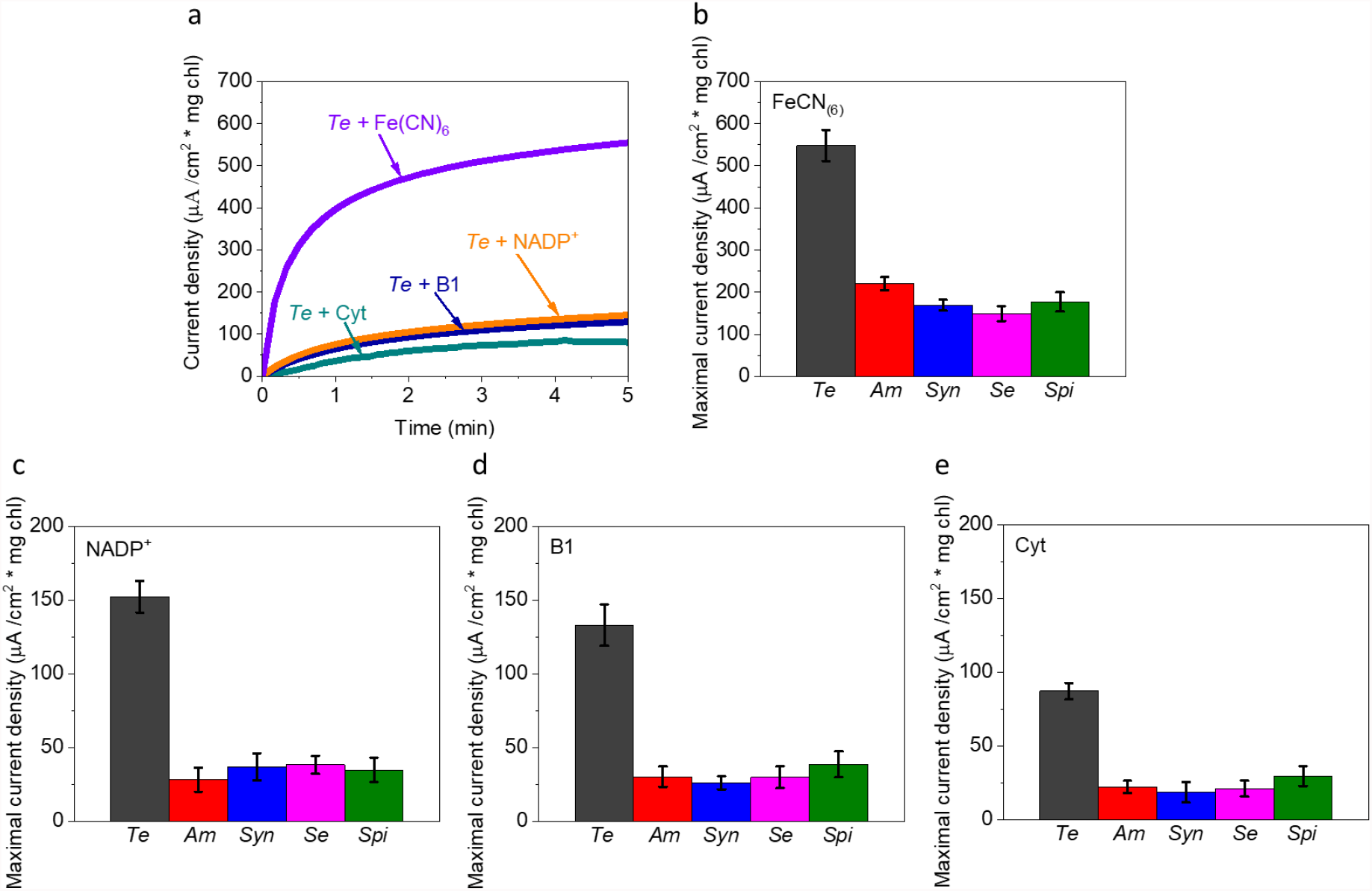
TE produces more current than other species in the presence of exogenous mediators. CA of the bacterial species *Te, Am, Syn, Se*, and *Spi* was measured in light with the addition of the exogenous electron mediators FeCN_(6)_, NADP^+^, B1, and Cyt. **a** Representative CA measurements of *Te* + FeCN_(6)_ (purple), NADP^+^ (orange), B1 (dark blue), and Cyt (azure). Maximal photocurrent densities of *Te, Am, Syn, Se*, and *Spi* with the addition of the exogenous electron mediators: **b** FeCN_(6)_, **c** NADP^+^, **d** B1, and **e** Cyt. The error bars represent the standard deviation over 3 independent measurements.

**Fig. 6.**
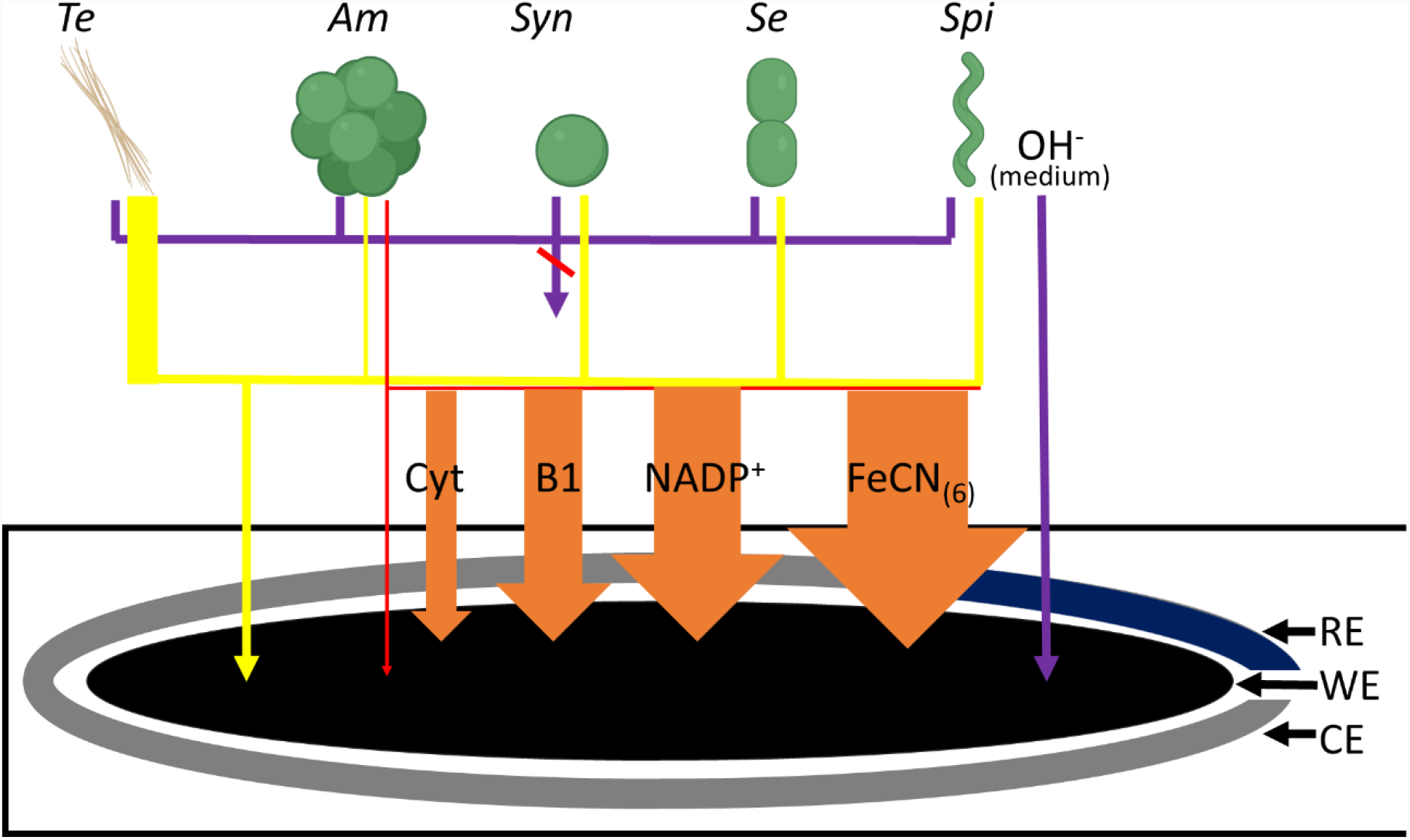
A schematic description of the factors that enhance the photocurrent in TE and other cyanobacterial-based BPECs. The cyanobacterial species *Te, Am, Syn, Se*, do not produce any significant current in dark. *Spi* does produce an electrical current in dark, however, this current does not originate from the cells It derives from the basic media of the cells (pH = 9) that consists of OH^-^ that can reduce the anode. Upon solar illumination, All 5 species can produce photocurrent. *Am* consist of both Chl a and d while the other species consist of Chl an only. Therefore, it can absorb a broader range of wavelengths and exploit it for photosynthesis and current production. As a marine organism, it can produce current in an electrolyte with a high salinity, which also contributes to the current enhancement. However, *Am* produces a photocurrent that is not higher than the freshwater cyanobacterial species *Syn*, and *Se*, probably because of its intolerance for high light. *Te* produces a photocurrent that is higher than the other species with and without the addition of an exogenous electron mediator. The photocurrent of all species can be significantly enhanced by the addition of exogenous electron mediators. The influence of the mediators on the photocurrent enhancement is Cyt < B1< NADP^+^ < FeCN_(6)_. The rectangular shape at the bottom represents the screen-printed electrode. A black circle represents the graphite WE, a silver arc represents the Pt CE, and a blue arc represents the Ag coated with AgCl RE. Labeled illustrations of cyanobacteria represent the 5 species. A purple arrow represents the current formation in the dark that is formed by OH^-^ in the media of *Spi*. Purple lines that connect into a purple arrow crossed by a red line represent the lack of current formation in dark. A thin red arrow represents the current that derives from the absorption of Chl d in *Am*, that is lower than the total current obtained under solar illumination. Yellow lines that connect into an arrow represent the photocurrent production. The thicker yellow line that connects with *Te* represents a photocurrent production that is greater than the other species. Labeled orange arrow with increasing widths represents the proportion of the photocurrent enhancement by the addition of the mentioned electron mediators.

## Conclusions

In this work, we show for the first time that photocurrent production can be generated from *Te* in a BPEC. This electricity generation is significantly greater than for other cyanobacterial species while using them in BPECs with and with the addition of exogenous electron mediators. We suggest that the high abundance of *Te* in nature and its high ability to produce photocurrent makes it an optimal species to be used in future clean energy technologies that are based on electricity production from cyanobacteria.

## Materials and Methods

### Chemicals

All chemicals were purchased from Merck unless mentioned otherwise.

### Cyanobacterial cultivation

#### Species cultivation

*Synechocystis* sp. PCC6803 (*Syn*) and *Synechococcus elongatus* PCC 7942 (*Se*) were cultivated in BG11 medium (pH = 6.8). *Acaryochloris marina MBIC 11017* (*Am* was cultivated in MBG11^41^ media (a BG11 medium containing 30 g/L sea salts (Instant Ocean). *Spirulina (Spi)* was cultivated in BG11 medium (pH = 9). The cultivation of *Syn, Se, Am*, and *Spi* was conducted in a growth chamber at a light intensity of about 100 μE / m^2^ s, shaking at 100 rpm and 29C°.

#### Samples preparation

O.D measurements of the cells were done at 750 nm (Nanodrop 2000 UV-Vis spectrophotometer, Thermo Fisher Scientific). Cell count was done under the microscope using a hemocytometer grid. For the measurements in the BPEC, log phase grown cells (O.D_750nm_ = 0.6 – 1.0) were used. Total Chlorophyll concentration was determined by 90% methanol extraction followed by absorption measurements at 665 nm for chl a and 696 nm for chl d. Calculation of Chl concentration was performed as previously described. ^42,43^

#### Chronoamperometry measurements

Chronoamperometry measurements were done using Plamsens3 potentiostat (Palmsens) connected to screen-printed electrodes (110, Metrohm Dropsens). The WE has a surface area of 0.5 cm^2^ Illumination of 1.5 SUN (1500 W / m^2^, calibrated at the electrode surface height without samples) was done using a solar simulator (Abet). In all measurements, a bias potential of 0.5 was applied on the graphite anode (this bias value was chosen because it produced the maximal signal / noise in the CA measurements). The volume of the measured samples of Am, Syn, Se, and *Spi* was 100 µL that was pipetted directly onto the printed electrodes panel. For *Te* samples, 100 µL of their cultivation media was first dropped directly on the printed electrode, and the cells were added afterward as described in Fig. 1a. The round boundaries of the measuring area of the panel induced a similar drop shape in all measurements that did not leak to the sides by the surface tension of the liquid. The drop was not stirred and the cells typically do not sediment spontaneously over the short measurement duration of 5 min. As preparation for fluorescence measurements, 100 µL of bacterial cells were filtrated through a 0.22 µm filter membrane to produce the ECM solution. each sample was diluted with fresh BPEC buffer to a final volume of 2 mL.

#### Fluorescence measurements

All fluorescence measurements were done using a Fluorolog 3 fluorimeter (Horiba) with excitation and emission slits bands of 4 nm. Quantification of NAD(P)H concentrations were calculated based on NADH calibration curve which was based on increasing concentrations measured at (λ(ex) = 350 nm, λ(em) = 450 nm). The lines of diagonal spots that appear in all of the maps presented here and in the following figures result from the light scattering of the Xenon lamp.^44^

## Acknowledgments

Funding was provided by a “Nevet” grant from the Grand Technion Energy Program (GTEP) and a Technion VPR Berman Grant for Energy Research. Some of the results reported in this work were obtained using central facilities at the Technion’s Hydrogen Technologies Research Laboratory (HTRL) supported by the Nancy & Stephen Grand Technion Energy Program (GTEP), the ADELIS Foundation, and the Solar Fuels I-CORE. We thank Dr. Rachel Edreii for her technical support. Yaniv Shlosberg is supported by fellowships of the Nancy & Stephen Grand Technion Energy Program (GTEP) and by a Schulich Graduate fellowship.

## Author contributions

Yaniv Shlosberg, Dina Spungin, Ilana Berman-Frank, and Noam Adir conceived the idea. Yaniv Shlosberg, Dina Spungin, and Noam Adir designed the experiments. Yaniv Shlosberg performed the main experiments. Dina Spungin and Neta Holland assisted in performing the experiments. Yaniv Shlosberg, Dina Spungin, Neta holland, Gadi Schuster, Ilana Berman-Frank, and Noam Adir wrote the paper. Ilana Berman-Frank, and Noam Adir supervised the entire research.

## Declaration of interests

The authors declare no competing interests.

## Supporting Information

**Fig. S1.**
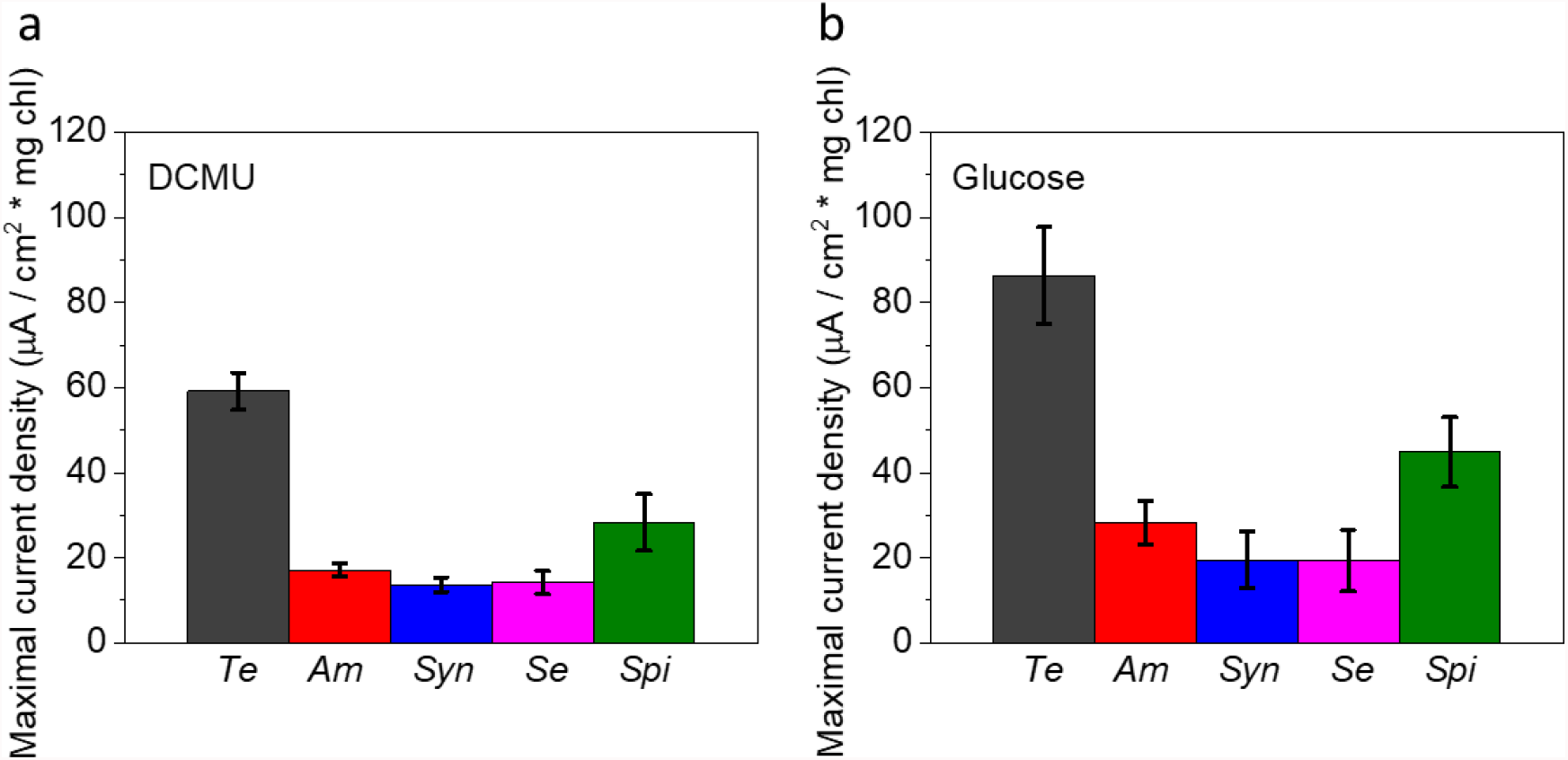
Addition of DCMU and Glucose enhances the photocurrent production in 5 cyanobacterial species. CA of the 5 cyanobacterial species *Te, Am, Syn, Se*, and *Spi* with addition of DCMU or Glucose were measured in light for 5 min. **a** Maximal photocurrent density of the 5 cyanobacterial species + DCMU. **b** Maximal photocurrent density of the 5 cyanobacterial species + Glucose. The error bars represent the standard deviation over 3 independent repetitions.

## Notes

### Competing Interest Statement

The authors have declared no competing interest.

